# Self-supervised Vision Transformers for image-to-image labeling: a BiaPy solution to the LightMyCells Challenge

**DOI:** 10.1101/2024.04.22.590525

**Authors:** Daniel Franco-Barranco, Aitor González-Marfil, Ignacio Arganda-Carreras

## Abstract

Fluorescence microscopy plays a crucial role in cellular analysis but is often hindered by phototoxicity and limited spectral channels. Label-free transmitted light microscopy presents an attractive alternative, yet recovering fluorescence images from such inputs remains difficult. In this work, we address the Cell Painting problem within the LightMyCells challenge at the International Symposium on Biomedical Imaging (ISBI) 2024, aiming to predict optimally focused fluorescence images from label-free transmitted light inputs. Leveraging advancements self-supervised Vision Transformers, our method overcomes the constraints of scarce annotated biomedical data and fluorescence microscopy’s drawbacks. Four specialized models, each targeting a different organelle, are pretrained in a self-supervised manner to enhance model generalization. Our method, integrated within the open-source BiaPy library, contributes to the advancement of image-to-image deep-learning techniques in cellular analysis, offering a promising solution for robust and accurate fluorescence image prediction from label-free transmitted light inputs. Code and documentation can be found at https://github.com/danifranco/BiaPy and a custom tutorial to reproduce all results is available at https://biapy.readthedocs.io/en/latest/tutorials/image-to-image/lightmycells.html.

## 1. INTRODUCTION

In the contemporary landscape of drug discovery and cellular analysis, Cell Painting has emerged as a transformative highcontent assay, renowned for its capacity to comprehensively profile cellular morphology [4]. This methodology employs six generic fluorescent dyes across five imaging channels to capture a diverse array of cellular components, including the nucleus, endoplasmic reticulum, nucleoli, RNA, actin, Golgi, plasma membrane, and mitochondria [4]. However, despite its cost-effectiveness, Cell Painting is constrained by the limited number of imaging channels available to prevent spectral overlap, thus restricting the scope of morphological information that can be obtained [5].

In contrast, label-free transmitted light microscopy methods such as brightfield (BF), phase contrast (PC), and Differential Interference Contrast (DIC) offer a compelling alternative. These techniques are non-invasive and significantly mitigate the drawbacks commonly associated with fluorescence microscopy, such as phototoxicity and the perturbation of cellular processes [6]. Fluorescence microscopy, while providing specificity, can inadvertently induce phototoxic effects and cytotoxicity, as well as interfere with the molecular interactions of its targets [6]. The innovation of in silico recovery of fluorescence images from brightfield images marks a groundbreaking shift in cellular imaging [6, 7, 8]. This approach, driven by advancements in deep learning and computer vision, not only addresses the limitations of fluorescent labeling but also enhances the information available from brightfield images. Thus, these methods stand at the forefront of a transformative movement in microscopy, promising enhanced sustainability and depth in cellular analysis.

Deep learning’s effectiveness across diverse sectors is well-documented, as highlighted in studies like [9]. However, in the biomedical field, its efficacy is often hindered by scarce annotated data, leading to small datasets. This limitation typically results in models that are inadequately trained and prone to overfitting, thereby diminishing their real-world applicability. In response to this issue, self-supervised learning has emerged as a promising approach, involving the creation of auxiliary tasks for model pretraining. Completing these tasks allows the model to gain additional insights, which can then be applied to down-stream tasks, enhancing the model’s knowledge representation and, consequently, its generalization performance.

Vision Transformers (ViT) [10] have sparked the development of numerous models, excelling in pretext task learning for self-supervised pretraining across different applications [11, 12, 13, 2, 14]. In medical image analysis, UNETR [1], stands as the first methodology employing a ViT as encoder, combined with a convolutional neural network (CNN) that functions as decoder.

In this work, we introduce an approach for the Cell Painting problem presented in the LightMyCells challenge at ISBI 2024. Our method aims to predict optimally-focused output images of various fluorescently labeled organelles from labelfree transmitted light input images. We leverage four specialized UNETR-like [1] models, each dedicated to predicting a specific organelle, and pretrained in a self-supervised manner. Our approach builds upon BiaPy, our open-source library for building deep-learning based bioimage analysis pipelines [3].

## 2. METHOD

### 2.1 Challenge Dataset

The dataset consists of previously unpublished 5D images from various studies, acquisition conditions, tissues, microscopes, resolutions, image size, and value ranges. The aim of the challenge is to predict different channels based on another, maintaining the time and channel axis constant (*dimension* = 1). The third axis represents focus, with ground truth extracted using the optimal focus algorithm [15]. Furthermore, the challenge aims to identify organelles with the best focus across different focal points. This yields approximately 57, 000 2D images, mostly sized at 2048 *×* 2048px, with 95% allocated for training. The remaining data is divided between preliminary and final test phases, with 10% and 90% of the remaining data, respectively.

Combining the total number of actin, nucleus, mitochondria and tubulin channels yields a sum of almost 4, 600 target images, which are distributed in 0.59%, 55.07%, 39.49% and 4.85% respectively. Input samples are divided into cases, each of them with a varying number of Z focus and output (target) images. Manual curation of the dataset was performed to identify and remove several fully black or blurred images.

### 2.2 Organelle-specialized 2D UNETR approach

The base model of our approach is a modified UNETR [1] within an image-to-image workflow developed using BiaPy [3]. Specifically, we deploy four distinct models, each dedicated to identifying a specific organelle and trained independently, as illustrated in Fig. 1.

**Fig. 1:**
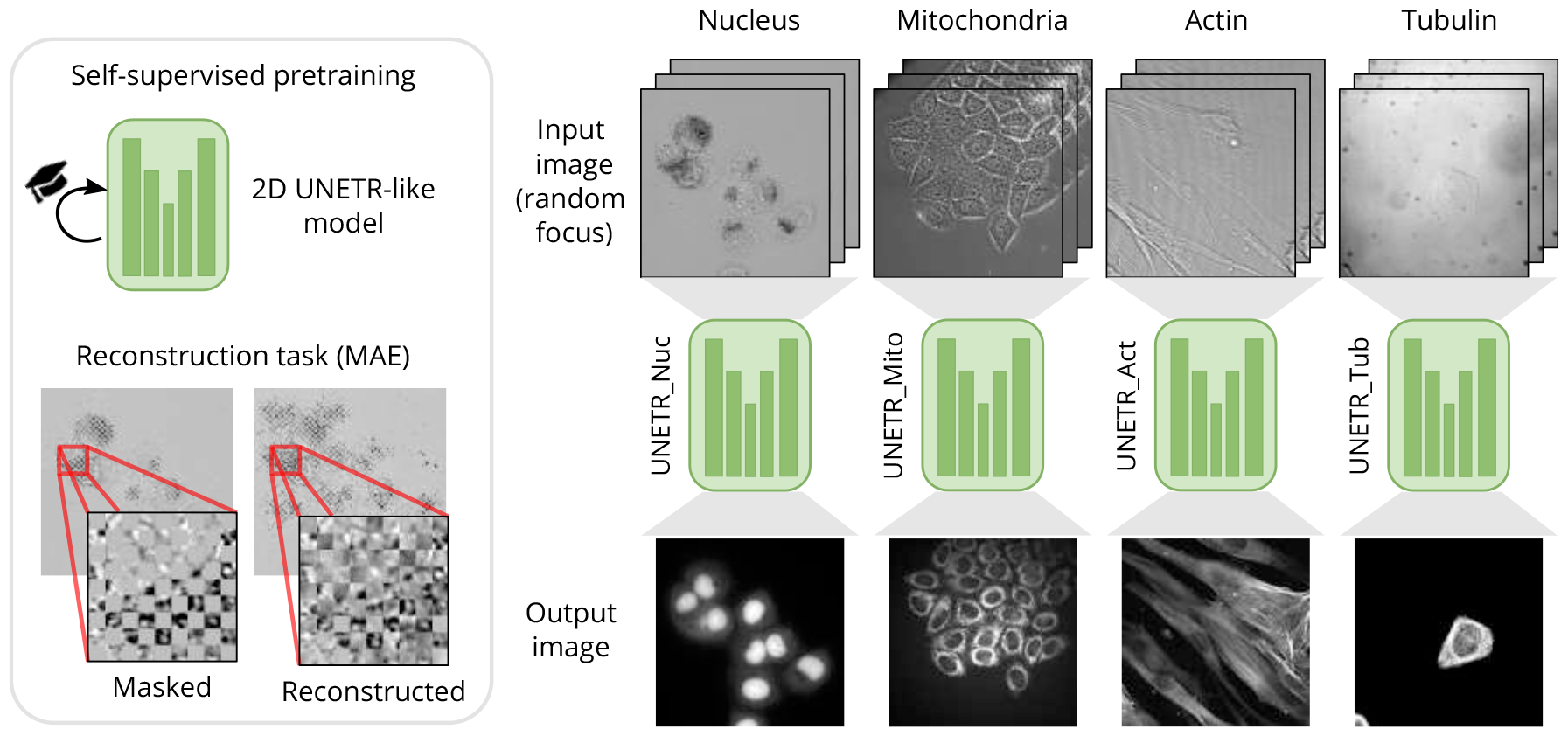
Schematic representation of our organelle-specialized 2D UNETR approach. The base model is a modified UNETR architecture [1] pretrained using MAE [2]. Then, four specialized models are fine-tuned independently for identifying specific organelles using a BiaPy [3] image-to-image workflow with heavy data augmentation.

#### Model specifications

We modified the original UNETR architecture to suit the challenge data. Namely, we adapted it to 2D, with an embedding size of 768, 12 layers, 12 heads and a token size of 32. Our UNETR decoder uses 32 filters on the first level (doubled on each level) and a kernel size of 9^1^ for each convolution.

#### Self-supervised pretraining

We pretrained the model encoder using all available images following the approach proposed in [2]. We employed a grid-masking occlusion instead of random occlusion. The research in [16] indicates that minimal occlusion is enough for a robust pretraining with a token size of 32 (50% of the image is masked in our case). Moreover, the images of this challenge are not object-centric [17]. Therefore, applying random masking often results in the removal of substantial information. This can lead to the model being unable to reconstruct the image effectively, forcing it to produce the mean value of each image. We followed the pretraining configuration of [2] but used a batch size of 4. Pretraining was conducted once, and weights were reused in all of the organelle-specialized models.

#### Training with random out-of-focus planes

The challenge dataset often includes multiple versions of the same sample captured at different out-of-focus planes. These images serve as different augmented versions of the sample. Therefore, we developed a custom data loader to randomly select one of these out-of-focus versions for each batch.

#### Heavy data augmentation

To enhance the models’ generalization capability, we applied random angle rotations, vertical and horizontal flips, elastic transformations, contrast and brightness changes, and Gridmask [18].

#### Implementation details

Our approach is fully integrated within the BiaPy library [3], using Pytorch version 2.2. We use an input size of 1024 *×* 1024, with mirroring employed to extend the image shape when it is smaller than the patch size. We set aside 10% of the training samples for validation, where all out-of-focus planes are used to assess the model’s performance. Mean squared error (MSE) was used as loss function, with a learning rate of 0.0001, employing a cosinedecay scheduler with warm-up [19], and ADAMW optimizer. All models were trained for 2500 epochs using eight A100 82GB GPUs with a patience of 250.

### 2.3 Exploration of training configurations and models

To investigate the relevance of each component in our approach, we perform a training configuration exploration of our two tested models: a CNN model (in Table 1a) and our final 2D UNETR-like [1] model (in Table 1b), that we used as baseline architectures. We measure the loss and peak signal-to-noise ratio (PSNR) between the ground truth and the prediction for the validation data used across the experiments. We compare six ablated versions with incremental changes (all applied except the one marked with * that employed “Heavy DA” in CNN exploration table). The initial CNN model was a five-level Residual U-Net model [20] using a patch size of 512 *×* 512, 32 feature maps in the first level of the network that are doubled on each level, batch normalization, a kernel size of 5, ADAMW optimizer with a learning rate base of 1.*E −* 4 (with cosine-decay scheduler with warm-up [19]) to minimize L1 loss, and 90º rotations, horizontal and vertical flips and brightness changes as data augmentation. We normalized all data (input and output) sample by sample, by subtracting the mean and dividing by the standard deviation. Both values are later used to denormalize the data before submission. We employed Contrast Limited Adaptive Histogram Equalization (CLAHE) to the data as pre-processing. For the baseline, we trained only with the best focus, meaning that for those images that had more than one focus we selected only the one with less blur^2^.

**Table 1:** Exploration of training configurations and models of our proposed approach for the LightMyCells challenge problem showing loss and PSNR metric measured on the validation data used across the experiments. From the top to the bottom, on each row, incremental modifications are applied based on the previous configuration (row marked with * in 1a not applied as decreases performance). The † represents the best configuration found using CNNs and then used with UNETR (as CNN experiments were done first). Best configurations for each organelle on each exploration table in gray . Best values in **bold**.

| CNN exploration |  |  |  |
| --- | --- | --- | --- |
| Target | Method | Loss↓ | PSNR↑ |
| Nucleus | Baseline (Residual U-Net) | 0.417 | 23.993 |
| Nucleus | Attention U-Net | 0.405 | 24.264 |
| Nucleus | Higher field of view (k=7) | 0.392 | 24.487 |
| Nucleus | Higher patch size: 1024x1024x1 | 0.344 | 25.210 |
| Nucleus | Not applying CLAHE | 0.276 | 29.324 |
| Nucleus | All image modalities | 0.258 | 30.620 |
| Nucleus | Higher field of view (k=9) | 0.251 | 30.765 |
| (*) Nucleus | Heavy DA | 0.298 | 29.823 |
| (†) Nucleus | All focuses picked at random | 0.252 | 28.219 |
| Mito. |  | 0.317 | 26.091 |
| Tubulin |  | 0.604 | 22.326 |
| Actin |  | 0.428 | 19.502 |

| UNETR exploration |  |  |  |
| --- | --- | --- | --- |
| Target | Method | Loss↓ | PSNR↑ |
| Nucleus | Baseline (UNETR) | 0.424 | 23.630 |
| Nucleus | Best CNN config (†) | 0.287 | 29.315 |
| Nucleus | MAE [2] pretraining | 0.259 | 30.217 |
| (*) Nucleus | Heavy DA | 0.228 | 30.976 |
| Nucleus | All focuses picked at random | 0.222 | 29.033 |
| Nucleus | MSE loss | 0.253 | 29.314 |
| Nucleus | Larger decoder | 0.243 | **29.546** |
| Mito. |  | 0.271 | **28.596** |
| Tubulin |  | 0.442 | **24.829** |
| Actin |  | 0.275 | **20.575** |

(b) Explored training configurations using UNETR-like model and their results on the validation sets.

In the initial weeks of the challenge, we encountered difficulties in achieving even minimal performance with a single model while simultaneously targeting multiple organelles. Consequently, we opted to concentrate initially on creating a robust starting point by targeting a single organelle using a single image modality. Our choice fell on the nucleus, with BF imaging. Upon implementing various modifications to our baseline model, we observed that the models failed to capture complete object shapes, resulting in patch/edge effects, even when predictions were made with overlap/padding. This led us to enhance the receptive field of the model by enlarging both the kernel size of all convolutions and the patch size, which yielded improved outcomes. A notable limitation of our derived solution was its lack of adaptability to different focal planes, as we were confined to using only the optimal focus when multiple options were available. To address this, upon achieving a satisfactory configuration, we incorporated the flexibility of selecting any random focus for each sample into our training regimen, increasing the model’s generalization capabilities.

Concurrently, we employed the optimal training configuration previously attained with our CNN (†) but used our UNETR-like [1] model instead. Although enhancements plateaued with the Attention U-Net following the application of additional data augmentation techniques on top of those already in use (as denoted by * in Table 1a), including random rotations and elastic transformations, the experiments with our UNETR-like model demonstrated superior generalization potential and a broader scope for improvement (indicated by * in Table 1b).

### 2.4 Results on LightMyCells challenge

The outcomes of our methodology are illustrated in Fig. 2. Nucleus and mitochondria are captured with great precision, in contrast to tubulin and actin, which present more challenges. This discrepancy is likely due to the limited volume of training samples available for these organelles (201 and 25 samples for tubulin and actin, respectively, whereas there nucleus and mitochondria are present in 2, 100 and 1, 455 samples, respectively).

**Fig. 2:**
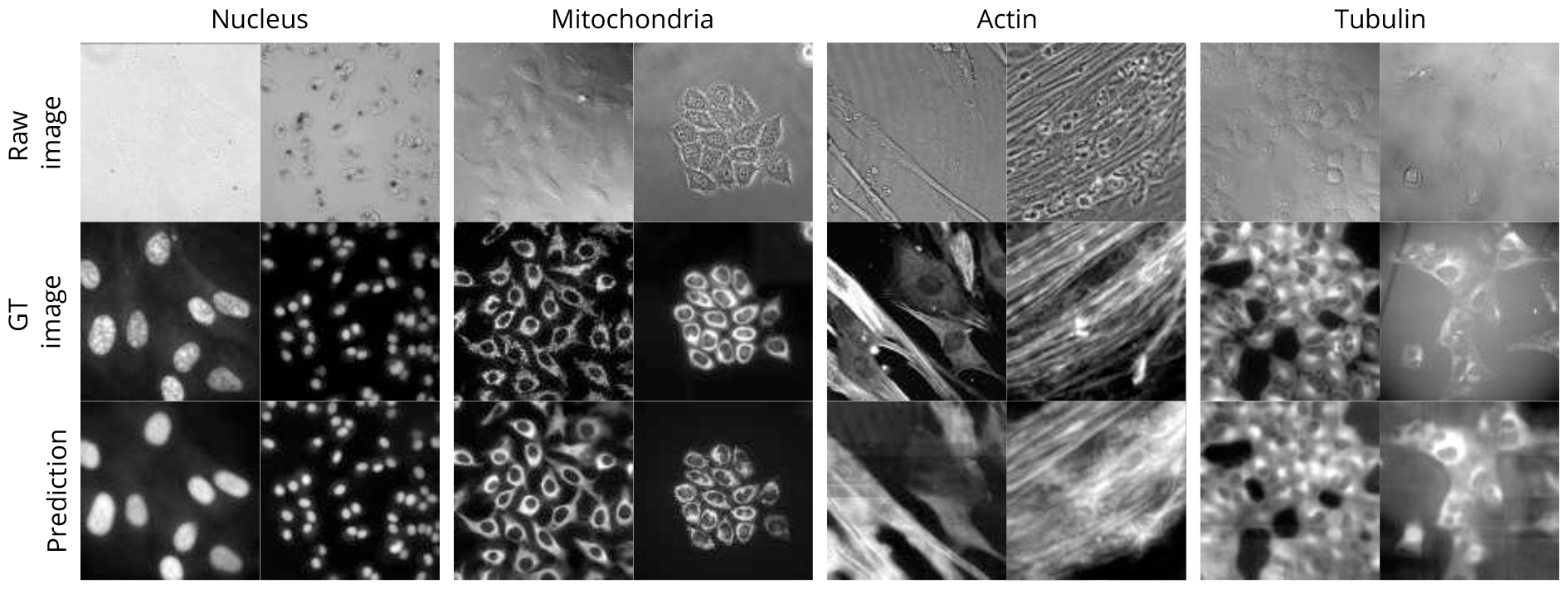
Results on the LightMyCells challenge of our approach.

## 3. CONCLUSION

In this paper, we present a vision transformer-based solution for the Cell Painting problem introduced in the LightMyCells challenge at ISBI 2024. Specifically, we propose four specialized UNETR-like models pretrained in a self-supervised manner, and each one dedicated to predicting a specific organelle. Experimental results demonstrate our proposed method is competitive without leveraging in the creation of any new architecture or loss, but using solid components and an organized experiment plan. All our experiments are fully reproducible as easy-to-use workflows of the open source BiaPy library.

## Acknowledgments

This work is partially supported by grant GIU23/022 funded by the University of the Basque Country (UPV/EHU), and grant PID2021-126701OB-I00 funded by the Ministerio de Ciencia, Innovación y Universidades, AEI, MCIN/AEI/10.13039/501100011033 and by “ERDF A way of making Europe”.

## Compliance with Ethical Standards

This work is a study for which no ethical approval was required.

## Footnotes

1 Excepting tubuling that best approach was obtained using 16 filters and a kernel size of 7

2 We estimate the strength of the blur with blur_effect() function of scikitimage [21]

